# T Cell Activation Triggers Reversible Inosine-5’-Monophosphate Dehydrogenase Assembly

**DOI:** 10.1101/315929

**Authors:** Krisna C. Duong-Ly, Yin-Ming Kuo, Matthew C. Johnson, Justin M. Kollman, Jonathan Soboloff, Glenn F. Rall, Andrew J. Andrews, Jeffrey R. Peterson

## Abstract

T cell-mediated adaptive immunity requires naïve, unstimulated T cells to transition from a quiescent metabolic state into a highly proliferative state upon T cell receptor engagement. This complex process depends on transcriptional changes mediated by Ca^2+^-dependent NFAT signaling, mTOR-mediated signaling and increased activity of the guanine nucleotide biosynthetic enzyme inosine-5’-monophosphate (IMP) dehydrogenase (IMPDH). Inhibitors of these pathways serve as potent immunosuppressants. Unexpectedly, we discovered that all three pathways converge to promote the assembly of IMPDH protein into micron-scale macromolecular filamentous structures in response to T cell activation. Assembly is post-transcriptionally controlled by mTOR and the Ca^2+^ influx regulator STIM1. Furthermore, IMPDH assembly and catalytic activity were negatively regulated by guanine nucleotide levels, suggesting a negative feedback loop that limits biosynthesis of guanine nucleotides. Filamentous IMPDH may be more resistant to this inhibition, facilitating accumulation of the higher GTP levels required for T cell proliferation.

**Abbreviations**
IMP: inosine-5’-monophosphate
IMPDH: inosine-5’-monophosphate dehydrogenase
LCMV: lymphocytic choriomeningitis virus
MPA: mycophenolic acid
mTOR: mechanistic target of rapamycin
NFAT: nuclear factor of activated T cells
STIM1: stromal interaction molecule 1
STIM2: stromal interaction molecule 2
TCR: T cell receptor

## Introduction

T cell receptor (TCR) stimulation initiates a complex series of events leading to T cell activation, proliferation, and differentiation. This core process of adaptive immunity is associated with dramatic metabolic alterations in T cells that are required for T cell proliferation (MacIver et al., 2013; Buck et al., 2015; Almeida et al., 2016; Bantug et al., 2018). The importance of these downstream signaling events is underscored by the fact that several immunosuppressive agents act by targeting these metabolic dependencies. For example, mechanistic target of rapamycin (mTOR) inhibitors and the inosine monophosphate dehydrogenase (IMPDH) inhibitor mycophenolic acid (MPA) are each used to suppress organ transplant rejection. The mechanisms by which these drugs effect immunosuppression are not fully understood (Allison and Eugui, 1996; Thomson et al., 2009; Nguyen le et al., 2015). However, restoring guanine nucleotide levels in MPA-treated T cells rescues their proliferation (Quemeneur et al., 2003), demonstrating that production of guanine nucleotides is the primary role of IMPDH in T cell activation. Intriguingly, mTOR also plays a key role in promoting guanine nucleotide synthesis (Ben-Sahra et al., 2016; Valvezan et al., 2017).

One of the earliest events of T cell activation is an increase in cytosolic Ca^2+^ content (Hogan et al., 2003; Gwack et al., 2007; Srikanth and Gwack, 2013). TCR-mediated Ca^2+^ signals are initiated by activation of phospholipase C leading to Ca^2+^ depletion in the endoplasmic reticulum (ER) that is sensed by stromal interaction molecule 1 (STIM1), a protein found in the ER membrane that serves an essential role in store-operated Ca^2+^ entry and is critical for sustained Ca^2+^ signals that drive the nuclear factor of activated T cells (NFAT) (Hogan et al., 2003; Gwack et al., 2007; Oh-Hora et al., 2008; Srikanth and Gwack, 2013). Hence, elevated cytosolic Ca^2+^ activates the serine/threonine phosphatase calcineurin, which dephosphorylates NFAT leading to nuclear import. Within the nucleus, NFAT is rapidly rephosphorylated, leading to its rapid export and highlighting the critical importance of sustained Ca^2+^ signals for T cell activation. Indeed, calcineurin inhibitors are potent immunosuppressants (Martinez-Martinez and Redondo, 2004), functioning primarily by blocking NFAT-mediated transcription. Importantly, the mTOR, IMPDH and STIM1/calcineurin pathways are thought to mediate the immune response via non-overlapping mechanisms. Here we present evidence that these pathways converge to promote the assembly of a macromolecular IMPDH-containing structure in a TCR-dependent process.

IMPDH is the rate-limiting enzyme in guanine nucleotide biosynthesis and is dramatically upregulated by T cell activation (Dayton et al., 1994). IMPDH has been recently reported to assemble into micron-scale structures in cultured cells, generally in response to pharmacological perturbations (Ji et al., 2006; Calise et al., 2014; Carcamo et al., 2014; Keppeke et al., 2015; Zhao et al., 2015; Calise et al., 2016). The physiological relevance of these filamentous assemblies is unknown, though they are reported to be catalytically active (Chang et al., 2015; Zhao et al., 2015; Anthony et al., 2017). We found that IMPDH assembles into filaments in primary T cells both *in vivo* and *ex vivo* in response to TCR engagement. Unexpectedly, we found that the assembly, but not the upregulated expression, of IMPDH was dependent on STIM1 and mTOR. This demonstrates that increased IMPDH expression is an early response to TCR stimulation but that IMPDH filament assembly is regulated by a post-translational mechanism mediated by STIM1 and mTOR. Thus, regulation of IMPDH is a common thread linking the pathways targeted by three major classes of immunosuppressive drugs. Our data suggests that IMPDH assembly may serve an essential function in T cell activation to support guanine nucleotide production.

## Results and Discussion

### TCR stimulation promotes IMPDH filament assembly in splenic T lymphocytes

We examined IMPDH expression and localization in primary T lymphocytes. T cells were isolated from the spleens of healthy mice and were activated using antibodies against the TCR co-receptors CD3ε and CD28 (Figure 1A). Strikingly, IMPDH assembled into linear assemblies and toroids in the vast majority of T cells within 24 hours (Figure 1A, B). Filament assembly was accompanied by a dramatic increase in IMPDH protein levels (Figure 1C). Together this demonstrates that increased IMPDH expression and filament assembly are direct downstream consequences of TCR activation and establish a system to analyze the regulation of these processes *in vitro*.

**Figure 1.**
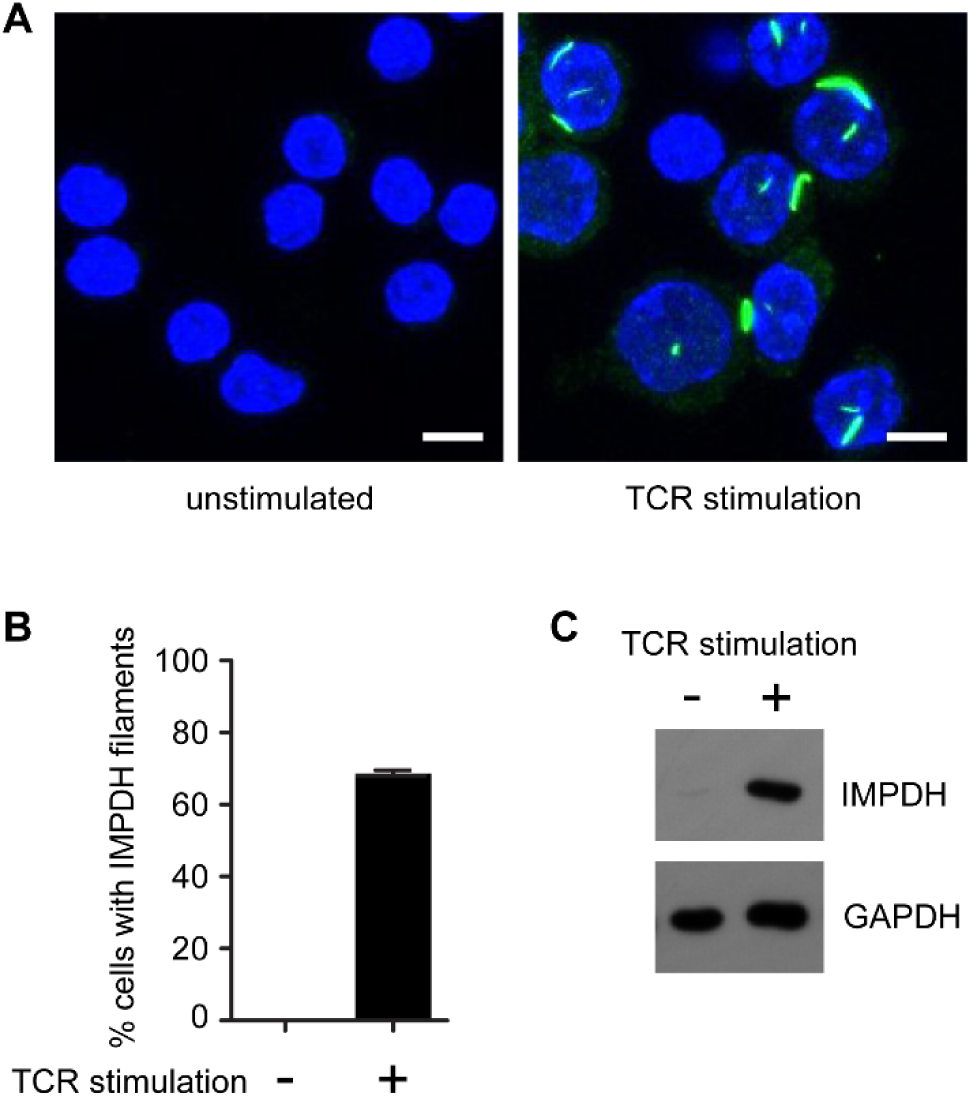
*Ex vivo* TCR stimulation promotes IMPDH protein expression and assembly into filaments. (A) Immunofluorescence images of splenic T cells that were either left unstimulated (left) or stimulated overnight with anti-CD3ε and anti-CD28 antibodies (right). IMPDH was stained using indirect immunofluorescence with an Alexa-488 conjugated secondary antibody (green) and nuclei were stained with DAPI (blue). Scale bar corresponds to 5 μm. (B) Quantification of the proportion of cells containing IMPDH filaments from three biological replicates consisting of 100 cells each. Error bars indicate standard error. (C) Western blot of IMPDH protein expression. GAPDH was used as a loading control. This blot is representative of eight biological replicates with similar results.

### STIM1 and mTOR regulate IMPDH filament assembly

To elucidate signaling mechanisms controlling IMPDH assembly upon T cell stimulation we first examined STIM1, an ER membrane protein that is a central mediator of calcium signaling (Hogan et al., 2003; Gwack et al., 2007; Srikanth and Gwack, 2013). To determine if STIM1 is important for filament formation, we compared IMPDH filament formation and expression in splenic T cells isolated from mice with a T cell-specific knockout of STIM1 (*STIM1^fl/fl^/Cd4-Cre*) and corresponding control mice (*STIM1^fl/fl^*) (Oh-Hora et al., 2008) and stimulated them *ex vivo*. There was a strong reduction of IMPDH filament formation in the STIM1-deficient cells compared to control (Figure 2A, B). Loss of STIM1 had only moderate effects on IMPDH protein expression, indicating that STIM1 is not primarily responsible for

**Figure 2.**
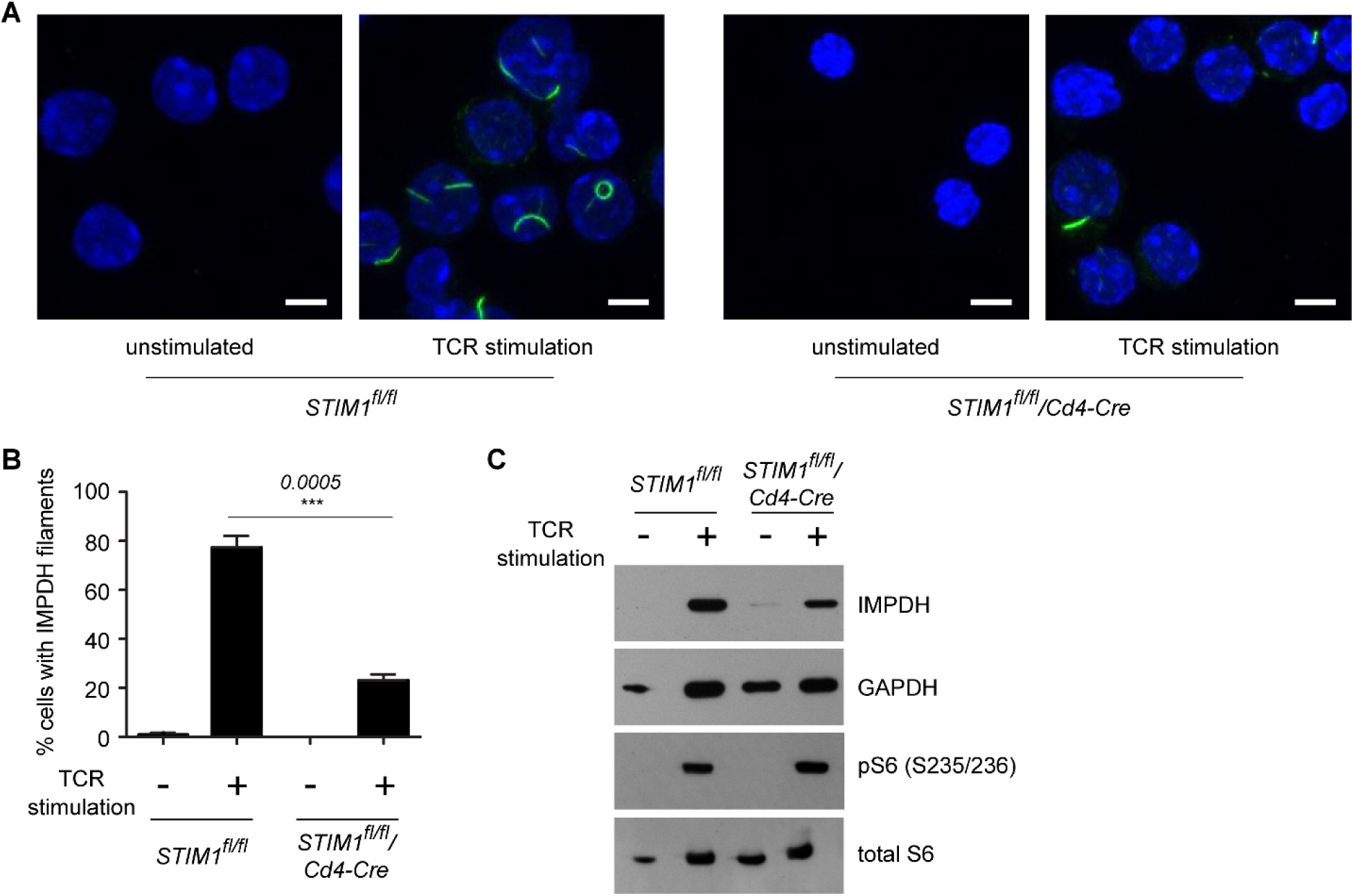
STIM1 regulates IMPDH filament assembly in an mTOR-independent manner. (A) Immunofluorescence images of splenic T cells isolated from *STIM1^fl/fl^* and *STIM1^fl/fl^/Cd4-Cre* mice that were either left unstimulated or stimulated *ex vivo* and immunostained as in Figure 1. Scale bars correspond to 5 μm. (B) Quantification of the proportion of T cells containing IMPDH filaments from three biological replicates consisting of 50–100 cells each. Error bars indicate standard error. The p-value for comparison of stimulated *STIM1^fl/fl^* and stimulated *STIM1^fl/fl^/Cd4-Cre* cells was computed from an unpaired two-sided Student’s t-test and is indicated above the error bar. (C) Western blot of IMPDH and phospho-S6 (pS6) ribosomal protein (Ser235/236) expression as a measure of mTOR activity. This result is representative of three biological replicates with similar results.

IMPDH upregulation (Figure 2C). Furthermore this finding demonstrates that IMPDH assembly is not a simple consequence of IMPDH upregulation, implying that assembly relies on a post-translational, Ca^2+^-dependent regulatory mechanism. Interestingly, IMPDH expression and assembly occurred normally in T cells isolated from mice deficient in the closely-related gene *STIM2* (Supplemental Figure 1), demonstrating a critical role for STIM1 specifically in post-translational regulation of IMPDH filament assembly in activated T cells.

The mTOR signaling pathway is a master regulator of diverse metabolic pathways and is a key mediator of the metabolic alterations during T cell activation (Chi, 2012; MacIver et al., 2013). Recently, mTOR was shown to promote purine biosynthesis (Ben-Sahra et al., 2016), in part as a mechanism to support ribosomal biogenesis (Valvezan et al., 2017). Conversely, purine levels have been shown to regulate mTORC1 activity (Emmanuel et al., 2017; Hoxhaj et al.), highlighting an intimate relationship between mTOR and purine nucleotides. Furthermore, mTOR is reported to act downstream of store operated calcium entry to promote metabolic alterations required for T cell activation (Vaeth et al., 2017). We therefore asked whether mTOR activity was important for IMPDH filament assembly.

Because mTOR could play roles both early and late in T cell activation, we stimulated T cells overnight to establish IMPDH filaments and then utilized small-molecule inhibitors to acutely inhibit mTOR (Figure 3A). One hour treatment with the allosteric mTOR inhibitor rapamycin almost completely disassembled IMPDH filaments (Figure 3B, C). Similar results were obtained using additional rapamycin analogs everolimus and temsirolimus as well as the ATP-competitive mTOR inhibitor AZD8055 (Figure 3B, C). The rapid disassembly of IMPDH filaments by mTOR inhibitors suggests that continued mTOR signaling is required to maintain these filaments. Furthermore, IMPDH filament disassembly by rapamycin was not due to any changes in IMPDH protein expression (Figure 3D), supporting the existence of a post-translational regulatory mechanism dependent on mTOR.

**Figure 3.**
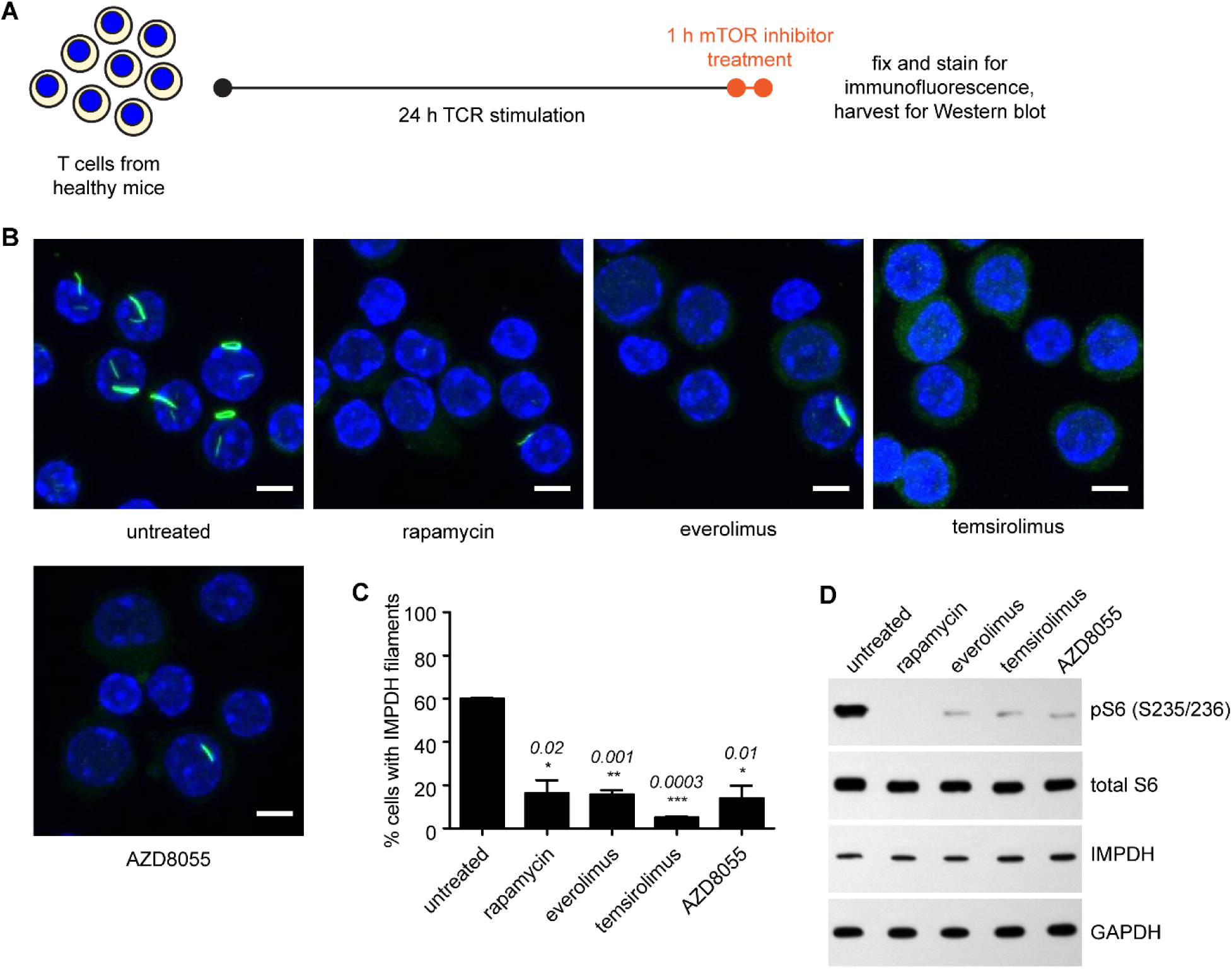
Continued mTOR activity is required for IMPDH filament maintenance. (A) Schematic of mTOR inhibitor treatment. (B) Immunofluorescence images of stimulated splenic T cells left untreated or treated with the indicated mTOR inhibitors. Cells were immunostained as in Figure 1. Scale bars correspond to 5 μm. (C) Quantification of the proportion of T cells containing IMPDH filaments from three biological replicates consisting of 100 cells each. Error bars indicate standard error. P-values for comparisons with control were computed using paired two-sided Student’s t-tests and are indicated above error bars. (D) Western blot of IMPDH and phospho-S6 ribosomal protein expression. The results are representative of three biological replicates.

Because mTOR has been proposed to act downstream of store operated calcium entry in T cells (Vaeth et al., 2017), we considered whether the failure of IMPDH to assemble in STIM1-deficient T cells was due to a loss of mTOR activity. However phosphorylation of the S6 ribosomal protein, a downstream effector of mTOR signaling, was similar in STIM1-deficient and control T cells (Figure 2C). This suggests that STIM1 and mTOR contribute independently to promoting IMPDH filament assembly downstream of TCR stimulation.

### IMPDH filaments in T cells are regulated by guanine nucleotide levels

IMPDH contains allosteric regulatory binding sites for certain guanine and adenine nucleotides. ATP promotes the assembly of recombinant IMPDH octamers into linear polymers (Labesse et al.; Anthony et al., 2017). These protofilaments are assumed to be a precursor in the assembly of the much larger IMPDH assemblies observed in cells. Within ATP-IMPDH filaments individual octamers can adopt either an expanded or a compressed conformation (Anthony et al., 2017). By contrast, binding of GDP or GTP to a partially overlapping site specifically promotes the collapsed form, which is catalytically inactive. Whether the collapsed form affects the stability of IMPDH polymers is unknown. Thus, relative cellular purine levels are thought to regulate both IMPDH polymer conformation and catalytic activity. To examine IMPDH filament responsiveness to nucleotides in T cells, we stimulated cells overnight to induce IMPDH assembly, and then supplemented the media with individual nucleosides for one hour. Following cellular uptake, nucleosides are readily converted to the corresponding mono-, di- and tri-phosphorylated nucleotides, bypassing *de novo* synthesis pathways (Figure 4A). Guanosine addition triggered the rapid disassembly of IMPDH filaments whereas filament number and morphology were unchanged in the presence of other nucleosides (Figure 4B, C). Furthermore, the loss of IMPDH filaments in the presence of guanosine was not associated with alterations in IMPDH protein levels (Figure 4D). Thus, guanosine and/or its downstream metabolites promote IMPDH filament disassembly.

**Figure 4.**
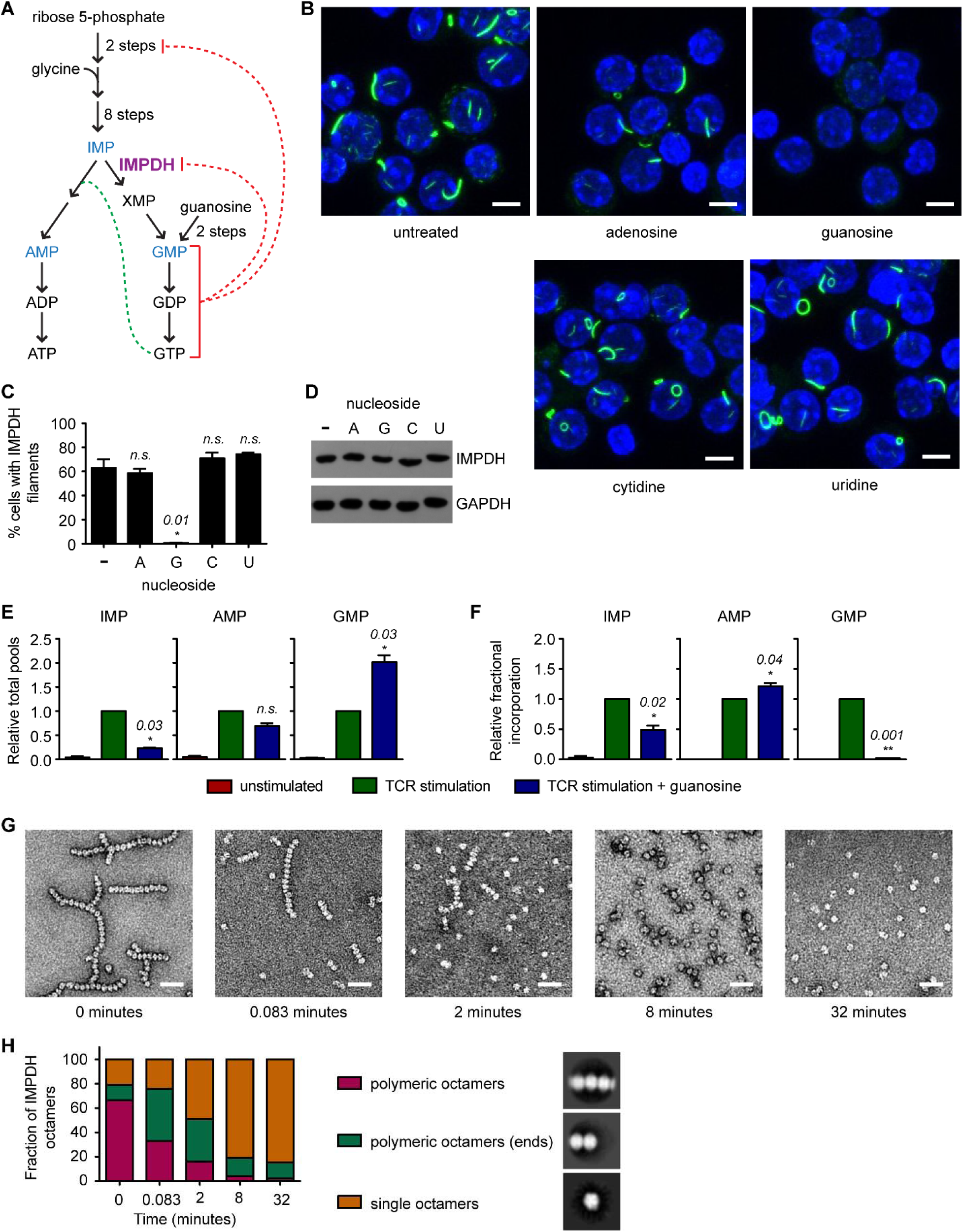
Guanosine specifically disassembles IMPDH filaments and inhibits IMPDH catalytic activity in activated T cells. (A) Schematic illustrating *de novo* purine biosynthesis. Positive and negative feedback regulation by guanine nucleotides is indicated by dashed green and red lines, respectively. (B) Immunofluorescence images of stimulated splenic T cells that were either treated with 200 μM of the indicated nucleoside for the last hour of stimulation or were left untreated. Cells were immunostained as in Figure 1. Scale bars correspond to 5 μm. (C) Quantification of the proportion of 100 T cells containing IMPDH filaments. Error bars indicate standard error. P-values for comparisons with control were computed as in Figure 3 and are indicated above error bars. (D) Representative western blot of IMPDH protein expression. (E) Total nucleotide pools in splenic T cells treated as indicated and subsequently incubated with [^13^C2,^15^N]-glycine for one hour prior to cell harvesting. Data represent three biological replicates and error bars correspond to standard error. P-values for comparisons with untreated stimulated T cells are given above error bars. (F) Relative fractional incorporation of [^13^C_2_,^15^N]-glycine for samples in (E). (G) Negative-stain electron microscopy images of 1 recombinant human IMPDH2 protein treated with 1 μM ATP at room temperature for 15 minutes to induce assembly. Subsequently, 1 mM GTP was added and samples were removed at the indicated times for analysis. Scale bar corresponds to 50 nm. (H) Quantification of data shown in (G). 500–1800 octamers were analyzed at each timepoint.

To investigate the effect of guanosine addition on *de novo* purine nucleotide biosynthesis, we monitored the incorporation of isotopically labeled glycine into purine nucleotides after guanosine treatment of activated splenic T cells. Glycine is an early precursor in *de novo* purine biosynthesis (Figure 4A) and is incorporated into the purine ring. T cells stimulated overnight or left unstimulated were incubated with [^13^C2,^15^N]-glycine for one hour and both total and [^13^C2,^15^N]-glycine-labeled IMP, AMP, and GMP were quantified by liquid chromatography/tandem mass spectrometry. IMP is the last common biosynthetic precursor of adenine and guanine nucleotides and AMP and GMP were chosen as representative nucleotides of each pathway.

TCR stimulation led to a dramatic upregulation of total IMP, AMP and GMP pools compared to unstimulated controls (Figure 4E), consistent with previous reports (Fairbanks et al., 1995). This was at least partially due to increased *de novo* purine biosynthesis since a corresponding increase was observed in the fraction of each nucleotide containing the [^13^C2,^15^N]-glycine-derived isotopomer in stimulated versus unstimulated cells (Figure 4F); however, it should be noted that stimulated cells do exhibit higher uptake of exogenous glycine (Supplemental Figure 2). Guanosine treatment led to a ~2-fold further increase in total GMP pool size (Figure 4E). Strikingly, this was associated with an almost complete inhibition of [^13^C2,^15^N]-glycine incorporation into GMP, demonstrating that although guanosine addition increased the total GMP pool, it suppressed *de novo* GMP synthesis (Figure 4F). These observations are best explained by the conversion of exogenous guanosine to form unlabeled GMP, GDP and GTP. GMP directly inhibits IMPDH catalytic activity competitively with the substrate IMP (Gilbert et al., 1979) while GDP and GTP inhibit IMPDH allosterically by promoting the catalytically inactive, collapsed conformation of IMPDH (Anthony et al., 2017; Buey et al.). Guanine nucleotides are also known to exert feedback control at two additional points in purine biosynthesis; GMP inhibits the synthesis of phosphoribosyl pyrophosphate in the rate-limiting first step of purine biosynthesis and GTP is a co-factor required by adenylosuccinate synthase activity to divert IMP toward adenine nucleotide biosynthesis (Figure 4A). Consistent with this, we found that guanosine addition decreased total IMP pools by half and suppressed incorporation of [^13^C2,^15^N]-glycine into IMP (Figure 4F). We also observed a slight, non-significant decrease in total AMP pool size, perhaps due to the decreased availability of IMP (Figure 4E). However, the incorporation of [^13^C2,^15^N]-glycine into AMP was increased (Figure 4F), suggesting that any IMP synthesized under conditions of excess guanosine is shunted to adenine nucleotide biosynthesis in the absence of IMPDH activity.

In conjunction with the immunofluorescence data, these results demonstrate for the first time that IMPDH filaments in a primary tissue are sensitive to alterations in guanine nucleotide levels. Increased guanine nucleotides are associated with the disassembly of IMPDH filaments and an inhibition of IMPDH catalytic activity. To test if these two events are mechanistically connected, we examined whether GTP destabilizes IMPDH polymers formed *in vitro*. Purified, recombinant IMPDH2 was incubated with ATP to promote assembly (Labesse et al., 2013; Anthony et al., 2017) and samples of the reaction were analyzed by negative stain electron microscopy before and after the addition of GTP for various times. GTP triggered disassembly of IMPDH polymers (Figure 4G, H), suggesting that the dissolution of IMPDH filaments in T cells treated with guanosine is mediated by the direct action of guanine nucleotides on IMPDH and is associated with the inhibition of IMPDH catalytic activity.

Our results strongly suggest that IMPDH filaments in stimulated T cells are catalytically active, consistent with prior observations that recombinant IMPDH polymers are catalytically active (Chang et al., 2015; Anthony et al., 2017). Excess guanine nucleotides suppress IMPDH assembly and activity, shunting IMP into adenosine nucleotide biosynthesis, presumably to restore balance to the adenine and guanine nucleotide pools. Since guanosine treatment rescues proliferation of T cells treated with the IMPDH inhibitor MPA (Quemeneur et al., 2003), guanine nucleotide production must be the primary role of IMPDH in supporting T cell activation. Our data suggest that filament assembly may support IMPDH activity in this context.

### Viral infection promotes IMPDH filament formation in splenic T lymphocytes

Finally, we investigated whether IMPDH filament assembly occurs *in vivo* in the context of natural T cell activation. Lymphocytic choriomeningitis virus (LCMV) is a mouse pathogen and a widely used model to study T cell activation and function. In LCMV-infected mice, T cells recognizing LCMV antigens become activated and undergo cellular proliferation. Following this expansion phase and resolution of infection, ~95% of activated T cells undergo apoptosis and the remaining surviving memory T cells confer protection against future LCMV infection (Murali-Krishna et al., 1998).

Mice were either left uninfected or infected with LCMV for 7 days, a time of peak anti-viral CD8^+^ T cell cytotoxicity (Hassett et al., 2000; Knipe and Howley, 2013), and splenic T cells were isolated. Immunostaining revealed robust assembly of IMPDH filaments (Figure 5A). By contrast, IMPDH filaments were observed only rarely in T cells from uninfected animals (Figure 5A). T cells from infected mice typically contained one or two IMPDH filaments, however some cells expressed a larger number of smaller IMPDH filaments. Despite only a subset of isolated T cells showing IMPDH assemblies, Western blotting revealed a three-fold increase in IMPDH protein levels in total splenic T cells from LCMV-challenged compared to control mice (Figure 5B). Following LCMV clearance, surviving CD69^+^ memory T cells lacked IMPDH filaments (Figure 5C), demonstrating transient assembly of IMPDH during initial activation that does not persist in the non-proliferating, metabolically quiescent memory cells.

**Figure 5.**
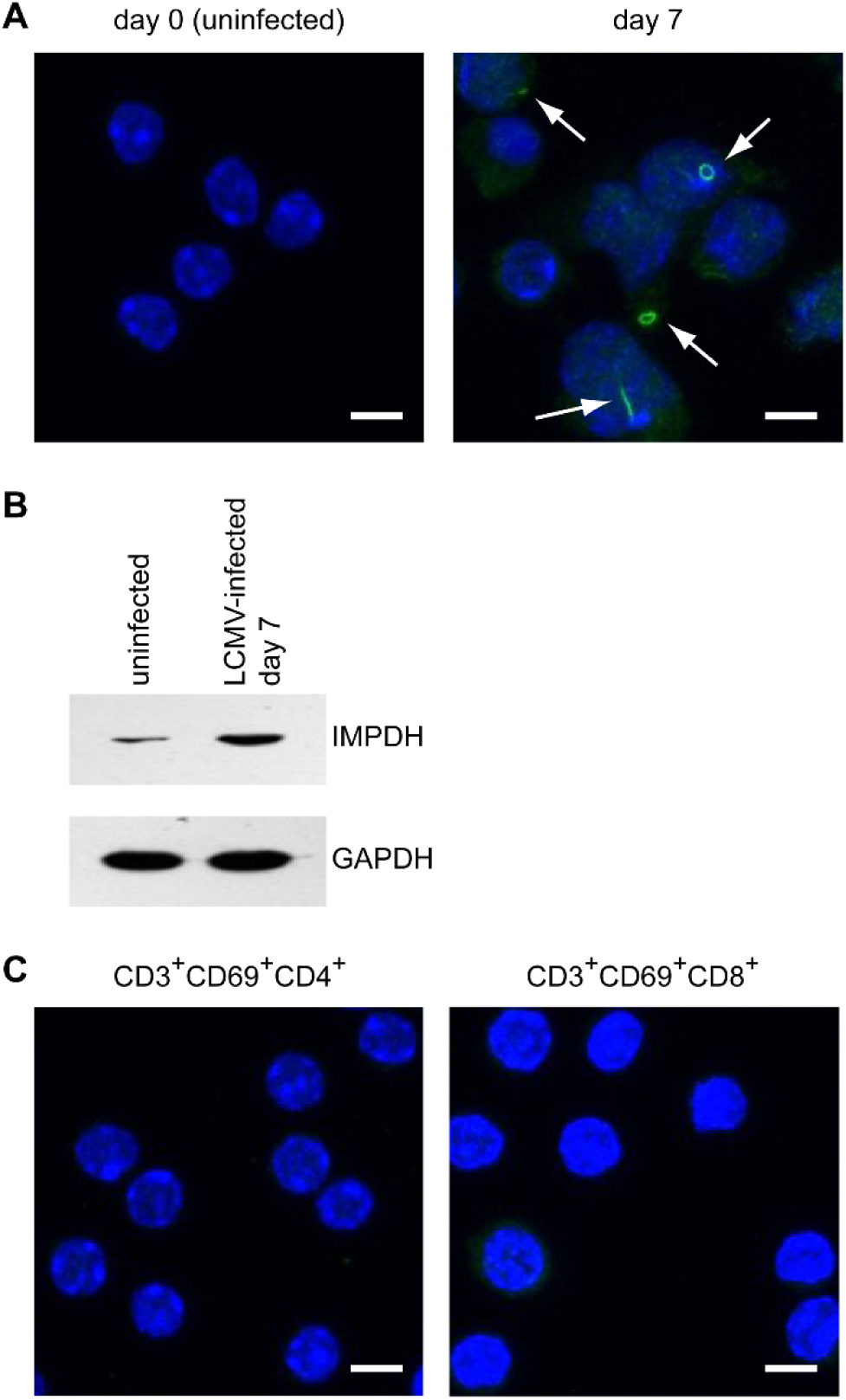
LCMV infection of mice promotes IMPDH filament assembly in vivo. (A) Immunocompetent C57BL/6 mice were infected intraperitoneally with 2×10^5^ plaque-forming units of LCMV-Armstrong virus. At the indicated times post-infection, red blood cells and T cells were depleted from splenocytes and IMPDH filament assembly was assessed by immunofluorescence at days 0, 7, and 30. White arrows highlight the location of IMPDH filaments. (B) Western blot of IMPDH protein expression of total splenic T cells isolated from uninfected mice or from mice 7 days after LCMV infection. (C) Immunofluorescence images of CD4^+^/CD69^+^ and CD8^+^/CD69^+^ memory T cells isolated from mice 30 days after LCMV infection. Cells were immunostained as in Figure 1. Scale bars correspond to 5 μm. These results are representative of three biological replicates.

Collectively, our results illustrate that IMPDH assembles into filamentous structures during T cell activation as a result of direct stimulation of the TCR. While IMPDH protein expression is highly upregulated at the protein level during this process, increased IMPDH expression is not sufficient to promote filament assembly; rather, IMPDH filament assembly is governed by posttranslational mechanisms including guanine nucleotide levels and downstream mediators of TCR signaling including mTOR and STIM1. More broadly, our findings introduce an experimentally tractable system to analyze IMPDH filament function and regulation in a physiologically relevant cell type.

Previous work demonstrated that polymerization *per se* does not enhance IMPDH catalytic activity (Anthony et al., 2017). However, our finding that guanosine treatment promotes both IMPDH filament dissolution and inhibition of IMPDH catalytic activity suggests the interesting possibility that filament assembly might serve to modulate IMPDH sensitivity to negative feedback regulation by guanine nucleotides. For example, IMPDH assemblies in T cells may stabilize the active, open conformation of IMPDH octamers, rendering them more resistant to octamer collapse and inactivation by GDP/GTP compared to free octamers. As a result, cells with filamentous IMPDH would be able to maintain higher steady state levels of guanine nucleotides needed to support T cell activation and proliferation. In contrast to this model, a prior study found that polymerization of recombinant IMPDH *in vitro* did not protect against GTP-mediated catalytic inhibition (Anthony et al., 2017). However, the linear polymers that were assembled *in vitro* may not reflect the activity of the larger IMPDH assemblies *in vivo*, which could also contain additional proteins or post-translational modifications that facilitate resistance to feedback inhibition.

Surprisingly, we find that three pathways essential for T cell activation, STIM1/calcineurin-mediated NFAT activation, signaling by mTOR, and IMPDH all converge to promote the assembly of IMPDH-containing filaments on TCR engagement. This observation may point to a critical role for filament assembly in T cell function. Indeed, we found that immune suppressing drugs that target mTOR disrupt IMPDH filament assembly. We expect that calcineurin-targeting agents would produce a similar effect based on the phenotype of STIM1-deficient T cells. In contrast, the IMPDH-targeted immunosuppressant MPA *promotes* IMPDH assembly in a variety of cell types (Ji et al., 2006; Calise et al., 2014; Carcamo et al., 2014; Keppeke et al., 2015). This could be a homeostatic response to reduced GTP levels in these cells or a direct consequence of MPA binding to IMPDH. Indeed, we found that MPA treatment also promoted IMPDH filament assembly even in unstimulated T cells without altering basal IMPDH protein levels (Supplemental Figure 3). This interesting observation demonstrates that increased IMPDH protein expression is not essential for filament assembly, consistent with a post-translational mechanism. More importantly, IMPDH assembly could serve as a simple pharmacodynamic biomarker in transplant patients treated with MPA.

## Materials and methods

### Animals

The Institutional Animal Care and Use Committees at Fox Chase Cancer Center and Temple University School of Medicine approved all animal procedures. Eight to 12 week old mice were used for all experiments. Wild-type C57BL/6 mice were either obtained from breeding colonies at the Fox Chase Cancer Center (FCCC) Laboratory Animal Facility or purchased from Taconic Biosciences. The generation of loxP-flanked STIM1 and STIM2 genes on the C57BL/6 background and the production of mice with T cells deficient in STIM1 and STIM2 has been previously described (Oh-Hora et al., 2008). For LCMV infection studies, immunocompetent C57BL/6 mice were infected with 2×10^5^ plaque forming units of LCMV-Armstrong as previously described (Matullo et al.) for either 7 or 30 days. All mice were euthanized by CO_2_ asphyxiation.

### Isolation of splenic T cells

Spleens were dissected from mice, crushed using the blunt end of a sterile syringe plunger, and passed through a 100 μm mesh filter incubating in T cell medium (RPMI-1640 supplemented with 10% heat-inactivated FBS, 100 U/mL penicillin, 100 μg/mL streptomycin, 2 mM L-glutamine, 1 mM sodium pyruvate, 50 μM β-mercaptoethanol, and non-essential amino acids [Gibco]). Splenocytes were harvested and resuspended in Ammonium chloride potassium (ACK) lysis buffer (155 mM Ammonium chloride, 10 mM Potassium bicarbonate, and 0.1 mM EDTA) to lyse red blood cells. Lymphocytes were harvested and subsequently resuspended in HBSS supplemented with 1% BSA. B cells were depleted by negative selection by binding to mouse CD45R (B220) microbeads (Miltenyi Biotec) and applying the cells to LS columns (Miltenyi Biotec) according to manufacturer’s instructions.

For CD69^+^ cell isolation, splenic T cells were isolated as described above and incubated with APC conjugated anti-CD3ε, BV605 conjugated anti-CD8a, APC-Cy7 conjugated anti-CD4, and PE conjugated anti-CD69 antibodies (BD Biosciences and Biolegend). Cell sorting was carried out on a 15-color BD FACSAria II instrument outfitted with 407, 488, 532, and 594 nm lasers. Data were collected with BD FACS Diva software. Single CD3^+^/CD69^+^/CD8^+^ and CD3^+^/CD69^+^/CD4^+^ cells represented 1.2 and 5.3% of the parental cell population, respectively.

### *Ex vivo* stimulation

Except where otherwise indicated, 24-well tissue culture plates were coated with 500 μL of a 10 μg/mL solution of anti-mouse CD3ε clone 145-2C11 (Biolegend) solution for 4 hours at 37°C. Coating solution was removed and wells were washed with T cell medium to remove excess anti-CD3ε. 1.5×10^6^ cells in 1 mL were applied to each well and anti-mouse CD28 clone 37.51 (Biolegend) was added to each well at a final concentration of 2 μg/mL. Stimulation was carried out for 24 hours in a humidified incubator at 37°C, 5% CO_2_. If used, mTOR inhibitors or guanosine were added for the last hour of stimulation directly to the media. Guanosine was used at the indicated concentrations while rapamycin was used at 20 nM, everolimus was used at 1 μM, temsirolimus was used at 10 μM, and AZD 8055 was used at 100 nM.

### Immunofluorescence and quantification

Cells were transferred to 8-well Lab-Tek II CC2 chamber slides (Thermo Scientific) and incubated at 37°C, 5% CO_2_ for 1 hour to allow cells to adhere to the surface. Cells were then fixed in 10% formalin diluted into PBS. Cells were subsequently washed with PBS, permeabilized with 0.5% Triton in PBS, and washed with 0.1% Triton. Non-specific binding was blocked with phosphate-buffered saline (PBS) supplemented with 0.1% Triton X-100, 2% bovine serum albumin (BSA), and 0.1% sodium azide. Anti-IMPDH diluted in blocking buffer was used to stain cells. Alexa 488 conjugated goat anti-rabbit secondary antibody was used to detect primary antibody binding and DAPI was used to stain nuclei. Vectashield mounting medium (Vector) was applied to the cells and 24×50 mm coverglasses (Fisher Scientific) were mounted on slides and sealed using clear nail polish.

Imaging was performed using the 63x oil immersion objective of a Leica Microsystems TCS SP8 Advanced confocal laser scanning microscope. Images were acquired using Leica Application Suite Advanced Fluorescence software at 400 Hz with a zoom factor of 2.0. For each image, z-stacks of focal planes of 0.5 μm depth were collected and presented as maximum projections.

### Western blotting

Cells were washed with PBS, resuspended and lysed in 30 μL sample buffer (62.5 mM Tris pH 6.8, 2% sodium dodecyl sulfate, 10% glycerol, 100 mM dithiothreitol, and 0.15 mM bromophenol blue) per 1×10^6^ cells, sonicated in a water bath for 5 minutes, and heated at 95°C for 90 seconds. Samples were run on SDS-PAGE gels and then transferred to nitrocellulose membranes using the Amersham TE 70 semidry transfer apparatus (GE Healthcare). Nitrocellulose membranes were blocked in a solution of 5% nonfat dry milk in Tris-buffered saline with 0.5% Tween-20. Anti-IMPDH2 (Abcam ab129165), anti-GAPDH (Santa Cruz Biotechnology sc-32233), anti-phospho-S6 ribosomal protein (Ser 235/236) (Cell Signaling Technology #4858), and anti-S6 ribosomal protein (Cell Signaling Technology #2217) were used as primary antibodies. Either HRP conjugated goat-anti rabbit or goat-anti mouse IgG (ThermoFisher Scientific) were used as secondary antibodies. Blots were visualized using Western Blotting Luminol Reagent (Santa Cruz Biotechnology) and exposed to film in a dark room.

### Glycine incorporation assay and nucleotide measurements

Splenic T cells were either left unstimulated or stimulated *ex vivo* as above except that 10 cm tissue culture plates were coated with 10 mL of the anti-mouse CD3ε solution and 30×10^6^ cells in 20 mL were stimulated per plate. Cells were either left untreated or treated with the indicated concentrations of guanosine 20 hours after stimulation. Three hours later, [^13^C2,^15^N]-glycine was added at a concentration of 35 mg/L. One hour later, cells were harvested and washed twice with Dulbecco’s PBS.

Samples were processed as described previously (Laourdakis et al.). Briefly, cells were resuspended in 70 μL of a solution of 0.5 M perchloric acid, 200 μΜ [^13^C_9_, ^15^N_3_]-CTP, vortexed for 10 seconds, and incubated on ice for 20 minutes. Lysates were then neutralized with 7 μL 5 M potassium hydroxide, vortexed for 10 seconds, and incubated on ice for 20 minutes. Debris was harvested at 11,000xg for 10 minutes on a microcentrifuge. Soluble lysates were further filtered using Amicon Microcon 0.5 mL YM-100 centrifugal filters as per manufacturer’s instructions.

Measurement of nucleotides was carried out on an Acquity I Class (Waters) ultra-performance liquid chromatography (UPLC) instrument coupled to a Xevo TQ-S micro triple quadrupole mass spectrometer (Waters). Five μL of each sample was loaded onto an Acquity UPLC HSS T3 column (1.8 mm, 2.1 mm × 50 mm; Waters) with an in-line column filter. Quantitative analysis was performed using the multiple reaction monitoring mode and MassLynx v. 4.1 software.

### Glycine uptake assay

Splenic T cells were either left unstimulated or stimulated *ex vivo* as above. To 1.5×10^6^ cells in 1 mL of media, a mixture of 35 μg of cold glycine and 10 μCi of [2-^3^H]-glycine was added. After a 1 hour incubation, cells were harvested, washed two times with PBS, and lysed with 1% SDS. Glycine uptake was assessed using a liquid scintillation counter (Beckman).

### Negative stain electron microscopy (EM) of recombinant IMPDH2 and quantification

Recombinant human IMPDH2 protein was expressed and purified as previously described (Anthony et al., 2017). Briefly, IMPDH2 was cloned into pSMT3-Kan and the resultant plasmid was used to transform BL21(DE3) competent *E. coli* cells. Cells were grown in LB media at 37°C to an OD_600_ of 0.8 and then cooled on ice for 5 minutes. Protein expression was then induced with 1 mM IPTG for 4 hours at 30°C. Cells were harvested and resuspended in a buffer containing 50 mM potassium phosphate, 300 mM potassium chloride, 20 mM imidazole, 0.8 M urea and Benzonase nuclease (Sigma) at pH 8. Cells were lysed by homogenization using the Emulsiflex-C3 (Avestin). His-SUMO-tagged IMPDH was purified by nickel affinity chromatography using Ni-NTA Agarose (Qiagen) and an elution buffer containing 50 mM potassium phosphate, 300 mM potassium chloride, and 500 mM imidazole at pH 8.0. The His-SUMO tag was cleaved using 1 mg ULP1 protease (Mossessova and Lima, 2000) per 100 mg of protein overnight at 4°C. The next day, 1 mM dithiothreitol and 0.8 M urea were added and the protein was concentrated using a 30,000 MWCO filter (Millipore). IMPDH2 was purified by gel filtration chromatography using either a Superdex 200 or a Superose 6 gel filtration column in a buffer containing 50 mM Tris, 100 mM potassium chloride, and 1 mM dithiothreitol at pH 7.4. Purified IMPDH2 was stored at −80°C as one-time use aliquots.

Negative stain EM was carried out as previously described (Anthony et al., 2017). IMPDH was diluted to 1 μM in assembly buffer (50 mM Tris, 100 mM potassium chloride, 1 mM dithiothreitol, and 1 mM ATP at pH 7.4) and incubated for 15 minutes at room temperature. 1 mM GTP was then added. At the indicated times, samples were applied to glow-discharged continuous carbon film EM grids and negative-stained with 1% uranyl formate. Transmission EM was performed using a FEI Tecnai G2 Spirit at 120 kV and a Gatan Ultrascan 4000 CCD interfaced with the Leginon software package (Suloway et al., 2005). Image processing was performed using Appion and RELION software (Lander et al., 2009; Scheres, 2012). Particles were picked manually from 59 electron micrographs, and classified into sixteen classes using Relion two-dimensional classification. These sixteen classes were pooled into three qualitative morphological groupings, corresponding to octamers that were either polymerized, at the ends of polymers, or non-polymeric.

## Acknowledgements

We thank Alana O’Reilly for helpful comments on the manuscript. This work was supported by grants from the National Institutes of Health (GM083025 to J.R.P., T32 CA009035 to K.C.D., GM117907 to J.S., GM102503 to A.J.A., and GM118396 to J.M.K.), funds from the Pennsylvania Department of Health to J.R.P. and the F.M. Kirby Foundation to G.F.R., and a Fox Chase Cancer Center Board of Associates Postdoctoral Fellowship Award to K.C.D.

## Supplemental Figures

**Supplemental Figure 1.**
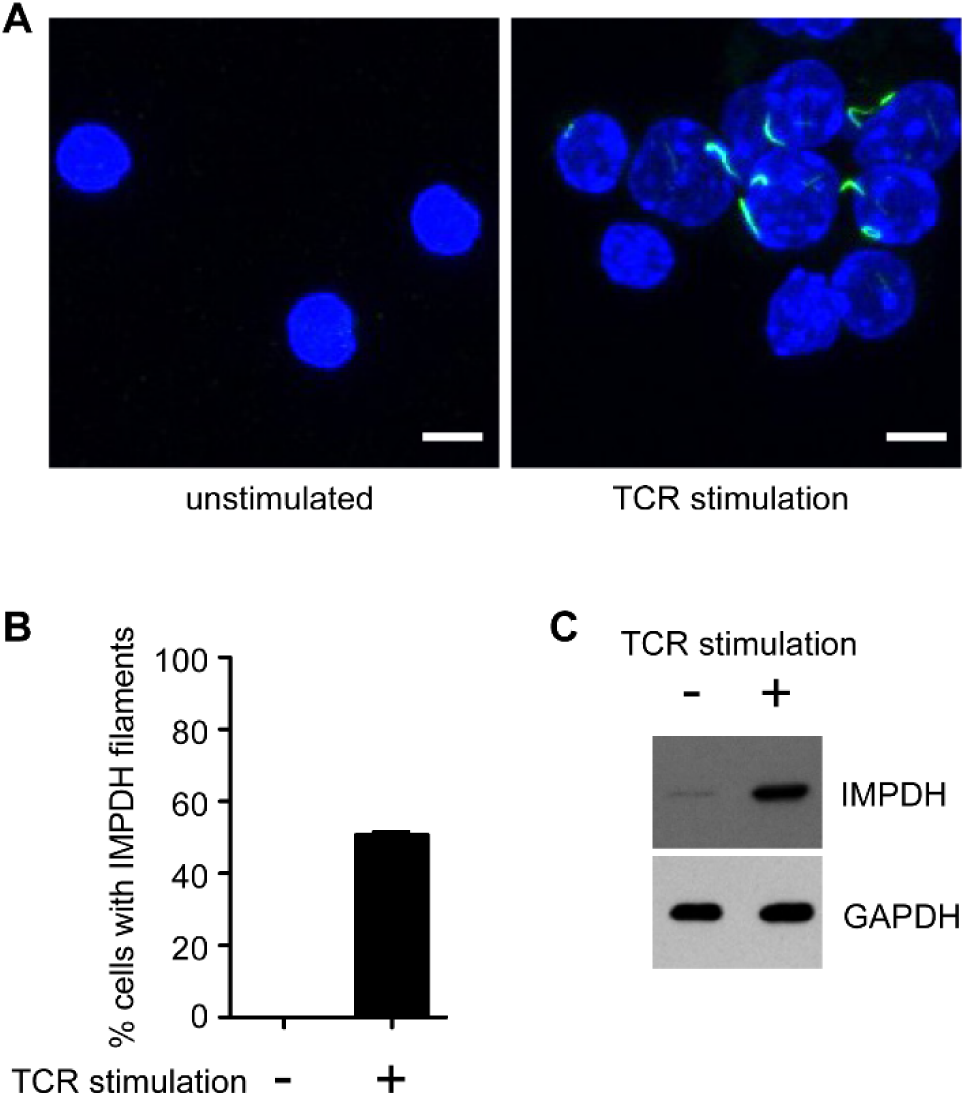
Deletion of STIM2 does not abrogate IMPDH filament assembly. (A) Immunofluorescence images of splenic T cells isolated from *STIM2^fl/fl^/Cd4-Cre* mice that were either left alone or stimulated *ex vivo* and immunostained as in Figure 1. Scale bars correspond to 5 μm. (B) Quantification of the proportion of cells containing IMPDH filaments from three biological replicates consisting of 100 cells each. Error bars indicate standard error. (C) Western blot of IMPDH protein expression. GAPDH was used as a loading control. This result is representative of three independent experiments.

**Supplemental Figure 2.**
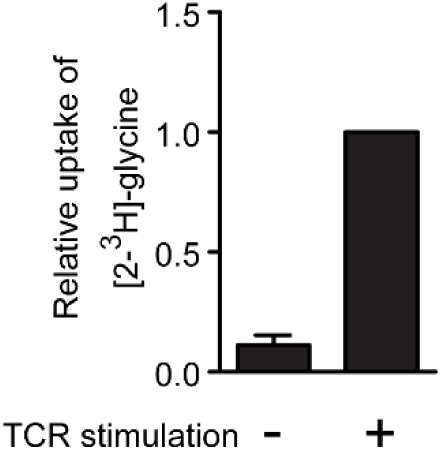
Uptake of exogenous glycine is increased upon TCR stimulation. Wild type splenic T cells were isolated and either stimulated or left alone (as in Figure 1) and then [2- ^3^H]-glycine was added for one hour. Cells were then washed to remove excess glycine remaining in the media and uptake was assessed using a liquid scintillation counter. Uptake values are plotted relative to that of stimulated T cells and error bars represent standard error. Two technical replicates were performed.

**Supplemental Figure 3.**
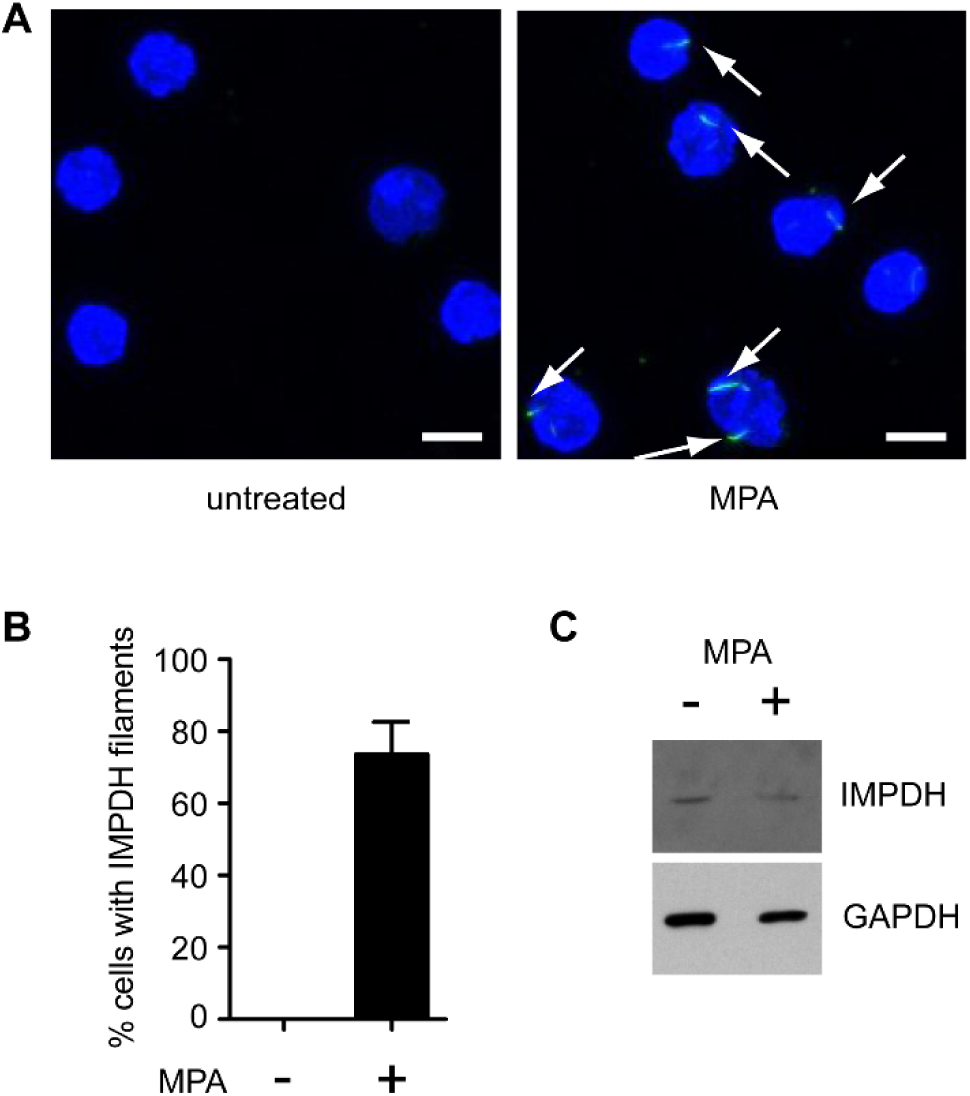
MPA treatment of unstimulated T cells promotes IMPDH filament assembly but not IMPDH expression. (A) Immunofluorescence images of unstimulated splenic T cells left untreated or treated with 10 μM MPA for 24 hours. Cells were immunostained as in Figure 1. Scale bars correspond to 5 μm. (B) Quantification of the proportion of T cells containing IMPDH filaments from three biological replicates consisting of 50–100 cells each. Error bars indicate standard error. (C) Western blot of IMPDH protein expression. GAPDH was used as a loading control. This result is representative of three independent experiments.

## References

Allison, A.C., and Eugui, E.M. (1996). Purine metabolism and immunosuppressive effects of mycophenolate mofetil (MMF). Clin Transplant 10, 77–84.

Almeida, L., Lochner, M., Berod, L., and Sparwasser, T. (2016). Metabolic pathways in T cell activation and lineage differentiation. Semin Immunol 28, 514–524.

Anthony, S.A., Burrell, A.L., Johnson, M.C., Duong-Ly, K.C., Kuo, Y.M., Simonet, J.C., Michener, P., Andrews, A., Kollman, J.M., and Peterson, J.R. (2017). Reconstituted IMPDH polymers accommodate both catalytically active and inactive conformations. Mol Biol Cell.

Bantug, G.R., Galluzzi, L., Kroemer, G., and Hess, C. (2018). The spectrum of T cell metabolism in health and disease. Nat Rev Immunol 18, 19–34.

Ben-Sahra, I., Hoxhaj, G., Ricoult, S.J.H., Asara, J.M., and Manning, B.D. (2016). mTORC1 induces purine synthesis through control of the mitochondrial tetrahydrofolate cycle. Science 351, 728–733.

Buck, M.D., O'Sullivan, D., and Pearce, E.L. (2015). T cell metabolism drives immunity. J Exp Med 212, 1345–1360.

Buey, R.M., Fernandez-Justel, D., Marcos-Alcalde, I., Winter, G., Gomez-Puertas, P., de Pereda, J.M., and Luis Revuelta, J. (2017). A nucleotide-controlled conformational switch modulates the activity of eukaryotic IMP dehydrogenases. Sci Rep 7, 2648.

Calise, S.J., Carcamo, W.C., Krueger, C., Yin, J.D., Purich, D.L., and Chan, E.K. (2014). Glutamine deprivation initiates reversible assembly of mammalian rods and rings. Cell Mol Life Sci 71, 2963–2973.

Calise, S.J., Purich, D.L., Nguyen, T., Saleem, D.A., Krueger, C., Yin, J.D., and Chan, E.K. (2016). 'Rod and ring' formation from IMP dehydrogenase is regulated through the one-carbon metabolic pathway. J Cell Sci 129, 3042–3052.

Carcamo, W.C., Calise, S.J., von Muhlen, C.A., Satoh, M., and Chan, E.K. (2014). Molecular cell biology and immunobiology of mammalian rod/ring structures. Int Rev Cell Mol Biol 308, 35–74.

Chang, C.C., Lin, W.C., Pai, L.M., Lee, H.S., Wu, S.C., Ding, S.T., Liu, J.L., and Sung, L.Y. (2015). Cytoophidium assembly reflects upregulation of IMPDH activity. J Cell Sci 128, 3550–3555.

Chi, H. (2012). Regulation and function of mTOR signalling in T cell fate decisions. Nat Rev Immunol 12, 325–338.

Dayton, J.S., Lindsten, T., Thompson, C.B., and Mitchell, B.S. (1994). Effects of human T lymphocyte activation on inosine monophosphate dehydrogenase expression. J Immunol 152, 984–991.

Emmanuel, N., Ragunathan, S., Shan, Q., Wang, F., Giannakou, A., Huser, N., Jin, G., Myers, J., Abraham, R.T., and Unsal-Kacmaz, K. (2017). Purine Nucleotide Availability Regulates mTORC1 Activity through the Rheb GTPase. Cell Rep 19, 2665–2680.

Fairbanks, L.D., Bofill, M., Ruckemann, K., and Simmonds, H.A. (1995). Importance of ribonucleotide availability to proliferating T-lymphocytes from healthy humans. Disproportionate expansion of pyrimidine pools and contrasting effects of de novo synthesis inhibitors. J Biol Chem 270, 29682–29689.

Gilbert, H.J., Lowe, C.R., and Drabble, W.T. (1979). Inosine 5'-monophosphate dehydrogenase of Escherichia coli. Purification by affinity chromatography, subunit structure and inhibition by guanosine 5'-monophosphate. Biochem J 183, 481–494.

Gwack, Y., Feske, S., Srikanth, S., Hogan, P.G., and Rao, A. (2007). Signalling to transcription: store-operated Ca2+ entry and NFAT activation in lymphocytes. Cell Calcium 42, 145–156.

Hassett, D.E., Slifka, M.K., Zhang, J., and Whitton, J.L. (2000). Direct ex vivo kinetic and phenotypic analyses of CD8(+) T-cell responses induced by DNA immunization. J Virol 74, 8286–8291.

Hogan, P.G., Chen, L., Nardone, J., and Rao, A. (2003). Transcriptional regulation by calcium, calcineurin, and NFAT. Genes Dev 17, 2205–2232.

Hoxhaj, G., Hughes-Hallett, J., Timson, R.C., Ilagan, E., Yuan, M., Asara, J.M., Ben-Sahra, I., and Manning, B.D. (2017). The mTORC1 Signaling Network Senses Changes in Cellular Purine Nucleotide Levels. Cell Rep 21, 1331–1346.

Ji, Y., Gu, J., Makhov, A.M., Griffith, J.D., and Mitchell, B.S. (2006). Regulation of the interaction of inosine monophosphate dehydrogenase with mycophenolic Acid by GTP. J Biol Chem 281, 206–212.

Keppeke, G.D., Calise, S.J., Chan, E.K., and Andrade, L.E. (2015). Assembly of IMPDH2-based, CTPS-based, and mixed rod/ring structures is dependent on cell type and conditions of induction. J Genet Genomics 42, 287–299.

Knipe, D.M., and Howley, P. (2013). Fields Virology.

Labesse, G., Alexandre, T., Vaupre, L., Salard-Arnaud, I., Him, J.L., Raynal, B., Bron, P., and Munier-Lehmann, H. (2013). MgATP regulates allostery and fiber formation in IMPDHs. Structure 21, 975–985.

Lander, G.C., Stagg, S.M., Voss, N.R., Cheng, A., Fellmann, D., Pulokas, J., Yoshioka, C., Irving, C., Mulder, A., Lau, P.W., Lyumkis, D., Potter, C.S., and Carragher, B. (2009). Appion: an integrated, database-driven pipeline to facilitate EM image processing. J Struct Biol 166, 95–102.

Laourdakis, C.D., Merino, E.F., Neilson, A.P., and Cassera, M.B. (2014). Comprehensive quantitative analysis of purines and pyrimidines in the human malaria parasite using ion-pairing ultra-performance liquid chromatography-mass spectrometry. J Chromatogr B Analyt Technol Biomed Life Sci 967, 127–133.

MacIver, N.J., Michalek, R.D., and Rathmell, J.C. (2013). Metabolic regulation of T lymphocytes. Annu Rev Immunol 31, 259–283.

Martinez-Martinez, S., and Redondo, J.M. (2004). Inhibitors of the calcineurin/NFAT pathway. Curr Med Chem 11, 997–1007.

Matullo, C.M., O'Regan, K.J., Curtis, M., and Rall, G.F. (2011). CNS recruitment of CD8+ T lymphocytes specific for a peripheral virus infection triggers neuropathogenesis during polymicrobial challenge. PLoS Pathog 7, e1002462.

Mossessova, E., and Lima, C.D. (2000). Ulp1-SUMO crystal structure and genetic analysis reveal conserved interactions and a regulatory element essential for cell growth in yeast. Mol Cell 5, 865–876.

Murali-Krishna, K., Altman, J.D., Suresh, M., Sourdive, D.J., Zajac, A.J., Miller, J.D., Slansky, J., and Ahmed, R. (1998). Counting antigen-specific CD8 T cells: a reevaluation of bystander activation during viral infection. Immunity 8, 177–187.

Nguyen le, X.T., Lee, Y., Urbani, L., Utz, P.J., Hamburger, A.W., Sunwoo, J.B., and Mitchell, B.S. (2015). Regulation of ribosomal RNA synthesis in T cells: requirement for GTP and Ebp1. Blood 125, 2519–2529.

Oh-Hora, M., Yamashita, M., Hogan, P.G., Sharma, S., Lamperti, E., Chung, W., Prakriya, M., Feske, S., and Rao, A. (2008). Dual functions for the endoplasmic reticulum calcium sensors STIM1 and STIM2 in T cell activation and tolerance. Nat Immunol 9, 432–443.

Quemeneur, L., Gerland, L.M., Flacher, M., Ffrench, M., Revillard, J.P., and Genestier, L. (2003). Differential control of cell cycle, proliferation, and survival of primary T lymphocytes by purine and pyrimidine nucleotides. J Immunol 170, 4986–4995.

Scheres, S.H. (2012). RELION: implementation of a Bayesian approach to cryo-EM structure determination. J Struct Biol 180, 519–530.

Srikanth, S., and Gwack, Y. (2013). Orai1-NFAT signalling pathway triggered by T cell receptor stimulation. Mol Cells 35, 182–194.

Suloway, C., Pulokas, J., Fellmann, D., Cheng, A., Guerra, F., Quispe, J., Stagg, S., Potter, C.S., and Carragher, B. (2005). Automated molecular microscopy: the new Leginon system. J Struct Biol 151, 41–60.

Thomson, A.W., Turnquist, H.R., and Raimondi, G. (2009). Immunoregulatory functions of mTOR inhibition. Nat Rev Immunol 9, 324–337.

Vaeth, M., Maus, M., Klein-Hessling, S., Freinkman, E., Yang, J., Eckstein, M., Cameron, S., Turvey, S.E., Serfling, E., Berberich-Siebelt, F., Possemato, R., and Feske, S. (2017). Store-Operated Ca(^2+^) Entry Controls Clonal Expansion of T Cells through Metabolic Reprogramming. Immunity 47, 664–679 e666.

Valvezan, A.J., Turner, M., Belaid, A., Lam, H.C., Miller, S.K., McNamara, M.C., Baglini, C., Housden, B.E., Perrimon, N., Kwiatkowski, D.J., Asara, J.M., Henske, E.P., and Manning, B.D. (2017). mTORC1 Couples Nucleotide Synthesis to Nucleotide Demand Resulting in a Targetable Metabolic Vulnerability. Cancer Cell 32, 624–638 e625.

Zhao, H., Chiaro, C.R., Zhang, L., Smith, P.B., Chan, C.Y., Pedley, A.M., Pugh, R.J., French, J.B., Patterson, A.D., and Benkovic, S.J. (2015). Quantitative analysis of purine nucleotides indicates that purinosomes increase de novo purine biosynthesis. J Biol Chem 290, 6705–6713.

